# Glycosylated proteins identified for the first time in the alga *Euglena gracilis*

**DOI:** 10.1101/2021.10.28.466288

**Authors:** Ellis O’Neill

## Abstract

Protein glycosylation, and in particular *N*-linked glycans, is a hall mark of Eukaryotic cells and has been well studied in mammalian cells and parasites. However, little research has been conducted to investigate the conservation and variation of protein glycosylation pathways in other eukaryotic organisms. *Euglena gracilis* is an industrially important microalga, used in the production of biofuels and nutritional supplements. It is evolutionarily highly divergent from green algae and more related to Kinetoplastid pathogens. It was recently shown that *E. gracilis* possesses the machinery for producing a range of protein glycosylations and make simple *N*-glycans, but the modified proteins were not identified. This study identifies the glycosylated proteins, including transporters, extra cellular proteases and those involved in cell surface signalling. Notably, many of the most highly expressed and glycosylated proteins are not related to any known sequences and are therefore likely to be involved in important novel functions in Euglena.

## 1. Introduction

Euglena are a class of mixotrophic protozoa that live in predominantly freshwater aquatic environments.^1^ Most possess a green secondary plastid derived by endosymbiosis of a chlorophyte algae,^2^ and there have been at least four endosymbiotic genome transfers, as well as significant horizontal gene transfer, during their evolutionary history.^3^ Uniquely among plastid containing cells, the chloroplast can be lost from photosynthetic Euglena without compromising their viability, due to duplication of all major pathways present in the chloroplast elsewhere in the cell.^4^ Euglenids are related to the well-known Kinetoplastid unicellular parasites *Trypanosoma* and *Leishmania*, as part of the phylum Euglenozoa.^5^ Euglena have been subject to scientific study for hundreds of years, but have recently become more intensely researched due to their considerable potential for biotechnological exploitation.^6^

*Euglena gracilis*, the most well characterised member of this group, has been studied for the production of vitamins A, C, E,^7^ essential amino acids, and polyunsaturated fatty acids.^8^ The storage polysaccharide, paramylon,^9^ makes up to 85% of algal dry weight when grown aerobically in light, whilst under anaerobic conditions wax esters can make up over 50% of the dry weight.^10^ These high value components have led to *E. gracilis* being cultivated as a food supplement.^11^ Recent work on the transcriptome and genome of *E. gracilis* has revealed the biosynthetic pathways for these valuable compounds.^12,13^

Euglena have been reported to have complex carbohydrates bound to their surface^14,15^ and lectin-and antibody-based profiling revealed a complex glycan surface, with some similarities to plant galactans and xylans.^16^ There are a wide range of carbohydrate active enzymes in the *E. gracilis* transcriptome, implying a capability for the synthesis of complex carbohydrates,^17^ and the cells contain a wide range of the sugar nucleotides needed as substrates for the synthesis of these polysaccharides.^16^ The exact nature of the complex surface carbohydrates in Euglena remains to be uncovered.

Protein glycosylation is a major post-translational modification in Eukaryotic organisms, stabilising surface proteins and providing specific intercellular interactions.^18^ *Euglena gracilis* expresses a range of enzymes necessary for the glycosylation of proteins: it has all of the genes necessary for the biosynthesis of GPI anchors, which anchor proteins into the phospholipid bilayer via a sugar-lipid tag, including the key transamidase for attaching the protein;^12^ there are three members of the GT41 family of glycosyltransferases, which transfer *N*-acetylglucosamine to serine and threonine residues of proteins in the cytosol;^17^ *N*-acetylglucosamine-1-phosphate transferase activity has been detected in membrane preparations of *E. gracilis* cells,^19^ likely involved in modifying proteins to target them to different subcellular compartments; sequences for all of the enzymes required for the synthesis of the highly conserved *N*-glycan precursor can be identified in the transcriptome, as well as three sequences for the transferases that transfer this pre-formed oligosaccharide to the target proteins.^12^ Together these results that Euglena encodes the ability to form complex posttranslational glycosylation of proteins. Protein *N*-glycan profiling of *E. gracilis* revealed that there was indeed protein glycosylation, mostly with high mannose type glycans with a small proportion modified with aminoethylphosphonate.^16^ No evidence was found for complex *N*-glycans or for *O*-linked glycans on *Euglena* proteins and the proteins carrying these modifications were not identified..

This study uses lectin mediated protein isolation and proteomic analysis to identify the proteins which are decorated with these glycans in order to understand the contribution of protein glycosylation to the Euglena proteome and inform future production of pharmaceutical proteins.

## 2. Results

Using standard proteomic techniques, the total proteome, glycan containing proteome and extracellular proteome were analysed from *Euglena gracilis* grown in a high yielding mixotrophic culture. It is notable that many of the most abundant proteins in all of the experimental samples in this study, as in previous work,^13^ are not linked to known sequences using BLAST. Many of those that do have known related sequences cannot be associated with GO terms or predicted functions, and together this indicates that some of the most highly abundant proteins in *E. gracilis* have no known function. It should also be noted that, due to limitations with the analytical techniques, the failure to detect a protein does not confirm its absence, but that it may not produce detectable peptides, be below the limit of detection or be masked by other, much more abundant, proteins.

The asparagine which is glycosylated can be identified by a mass deviation of 1 Dalton (Da) from the expected mass, caused by the cleavage of the *N*-glycan by PNGase-F treatment during the sample preparation. Peptides may not be detected in the modified form and the protein may be identified by other peptides, and so absence of this signal does not indicate absence of glycosylation of a protein. Only 88 of the 382 peptides annotated as containing this *N*-deamidation, appear to have the canonical NX(S/T) recognition signal for glycosylation. Whilst chemical deamidation of asparagine can occur giving rise to false identification of glycosylation sites,^20^ the high ratio of non-canonical sites in this data set raises serious concerns with the use of protein glycosylation site prediction tools that rely on this recognition motif for *Euglena* and indicates that there may be some other signal to target glycosylation in this organism. There are three sequences for the GT66 oligosaccharyltransferases that transfer the pre-formed glycan to the protein asparagine encoded in the Euglena transcriptome^12^ and it is possible that these have different specificities. Further experiments would be required to validate the true glycosylation sites.

### 2.1 Total proteins

1309 proteins were detected in all samples from the total proteome (see Supplementary Data), and of these 63% (836) have identified GO terms (see Figure 1), much higher than the 37% of the total transcripts which have GO terms mapped.^12^ Of the 130 proteins detected above the average, 30 do not have any BLAST hits, and 85 have GO terms identified. This indicates that the proteins that can be detected are less likely than those predicted from the transcriptome to have no known related sequences, possibly indicating the many of the predicted but unknown proteins are produced at a lower level or not at all. However, there are still many proteins that are unique to Euglena that are produced at relatively high levels and would repay further study.

**Figure 1:**
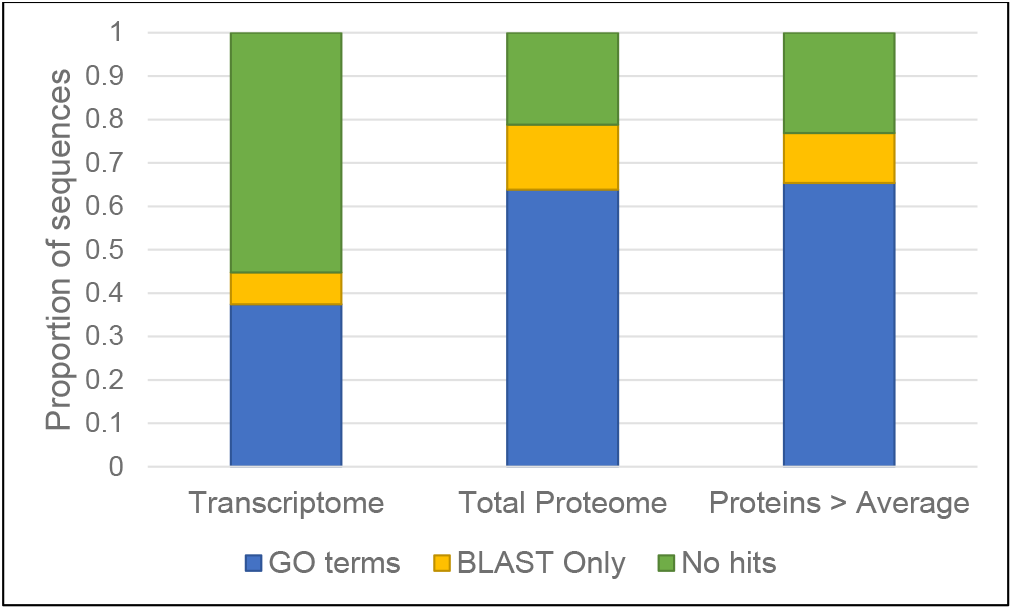
Euglena sequence identification. Proportion of sequences with identified GO terms and BLAST hits (blue), BLAST hits only (yellow) and neither (green) using Blast2GO, in the *E. gracilis* transcriptome,^12^ total proteome and proteins detected above average (this study).

In order to identify the likely subcellular location of these abundant proteins, protein targeting predictions were performed, using bioinformatic tools that have previously been used for Euglena proteins.^4^ Protein transport into *Euglena* chloroplasts occurs first via the secretory pathway and the Golgi apparatus using a secretion signal, followed by targeting to the chloroplast using a plastid targeting signal.^21^ Therefore, to confirm whether a protein was truly secreted or sent to the chloroplast, any predicted signal peptides were removed and the prediction repeated, revealing any masked plastid targeting signal. TargetP^22^ predicted that four of the 20 most abundant proteins are targeted to the mitochondria, two to the chloroplast and one secreted, whilst WoLF PSORT^23^ predicts six to be targeted to the chloroplast, four to the mitochondria and one secreted (see Table 1). These results suggest the chloroplast and mitochondria contain some of the most abundant proteins in the cell.

**Table 1:** The 20 most abundant proteins in the total Euglena proteome. The full list is available in the supplementary file. NA is sequences with no homologues identified by BLAST. Any secretory signal peptides identified were removed and the analysis repeated, with results shown in brackets. * indicates sequences that do not start with a M and so may be truncated sequences that do not contain the targeting sequence present in the protein.

| Sequence Name | Sequence Description | Average Intensity | StDev | TargetP prediction<sup>22</sup> | WoLF PSORT prediction<sup>23</sup> |
| --- | --- | --- | --- | --- | --- |
| 16406 | Glyceraldehyde 3-phosphate dehydrogenase | 17.44805 | 0.036709 | Other | Cytosol |
| 3371 | Stromal 70 kDa heat shock-related protein, chloroplastic-like | 13.07199 | 2.87894 | Chloroplast* | Chloroplast* |
| 12260 | Phosphoglycerate kinase | 9.826626 | 2.135535 | Other | Cytosol |
| 8709 | Oxygen-evolving enhancer protein 1, chloroplastic | 9.502916 | 1.305499 | Chloroplast | Chloroplast |
| 8430 | Putative mitochondrial heat-shock protein hsp70 | 9.346824 | 1.086321 | Other* | Cytosol* |
| 6665 | Phosphopyruvate hydratase | 8.758831 | 0.565104 | Other* | Cytosol* |
| 61646 | 60s acidic ribosomal protein p2 | 7.992886 | 0.289941 | Other* | Extracellular* (Chloroplast) |
| 4720 | NA | 7.732722 | 1.186309 | Other | Mitochondria |
| 25934 | Oxygen-evolving enhancer protein 3 | 7.419665 | 1.114722 | Signal Peptide (Signal peptide) | Chloroplast |
| 25617 | Electron transfer flavoprotein subunit alpha | 6.635908 | 1.382096 | Other* | Chloroplast* |
| 8433 | NA | 6.51502 | 0.534721 | Other | Nucleus |
| 8912 | Putative ATPase beta subunit | 6.505054 | 1.134881 | Mitochondria* | Mitochondria* |
| 12357 | Elongation factor 1-alpha | 5.914015 | 0.663056 | Other* | Cytosol* |
| 14010 | Beta-tubulin | 5.45031 | 0.352232 | Other* | Nucleus* |
| 7178 | Chaperonin HSP60, mitochondrial precursor | 4.972642 | 0.89093 | Mitochondria | Mitochondria |
| 23135 | NA | 4.893481 | 0.05412 | Other* | Cytosol* |
| 7258 | NA | 4.74644 | 0.335421 | Other | Nucleus |
| 8679 | Putative dihydrolipoamide dehydrogenase | 4.417853 | 0.969253 | Mitochondria | Chloroplast |
| 5458 | Molecular chaperone DnaK | 4.144819 | 2.138527 | Mitochondria | Mitochondria |
| 4275 | ATPase beta subunit | 3.973031 | 1.009266 | Other | Chloroplast |

### 2.2 ConA Glycoprotein Isolation

Concanavalin A (ConA) is a protein that specifically binds mannose, such as is found in simple *N*-glycans, and glucose which can be found on the termini of *N*-glycans. Using an immobilised ConA column to enrich for *N*-glycan displaying proteins, a total of 86 proteins were detected at a significantly higher rate than in the total proteome, and 50 of these were not detected in the total proteome at all (See Table 2). 36 of these ConA enriched proteins had BLAST matches and 30 mapped to GO terms. Six of these are likely to be involved in signalling, three in sugar metabolism, two in transport, and there are four likely proteases. There are four proteins that are linked to biosynthesis, two to redox balance, and 12 involved in core housekeeping roles, which would expect to be cytosolic and thus not glycosylated. 13 of the 86 proteins had an *N*-deamidation site detected in at least one of the samples. Of the proposed cytosolic housekeeping genes this modification was noted in: 7967, a trypanothione reductase that has a deamidation site in all ConA samples, as well as the single WGA sample in which it was detected; 5325, a small nucleolar ribonucleoprotein U3, with one *N*-deamidation site in just one ConA sample; 32750, a RNA scaffolding Sm-like protein, with deamidation in all WGA samples, although it was not detected significantly over the control in them, but not with no deamidation detected in any of the ConA samples. Only six of the ConA enriched proteins were predicted to be secreted, again highlighting the limitations of predicting protein targeting in protozoa.

**Table 2:**
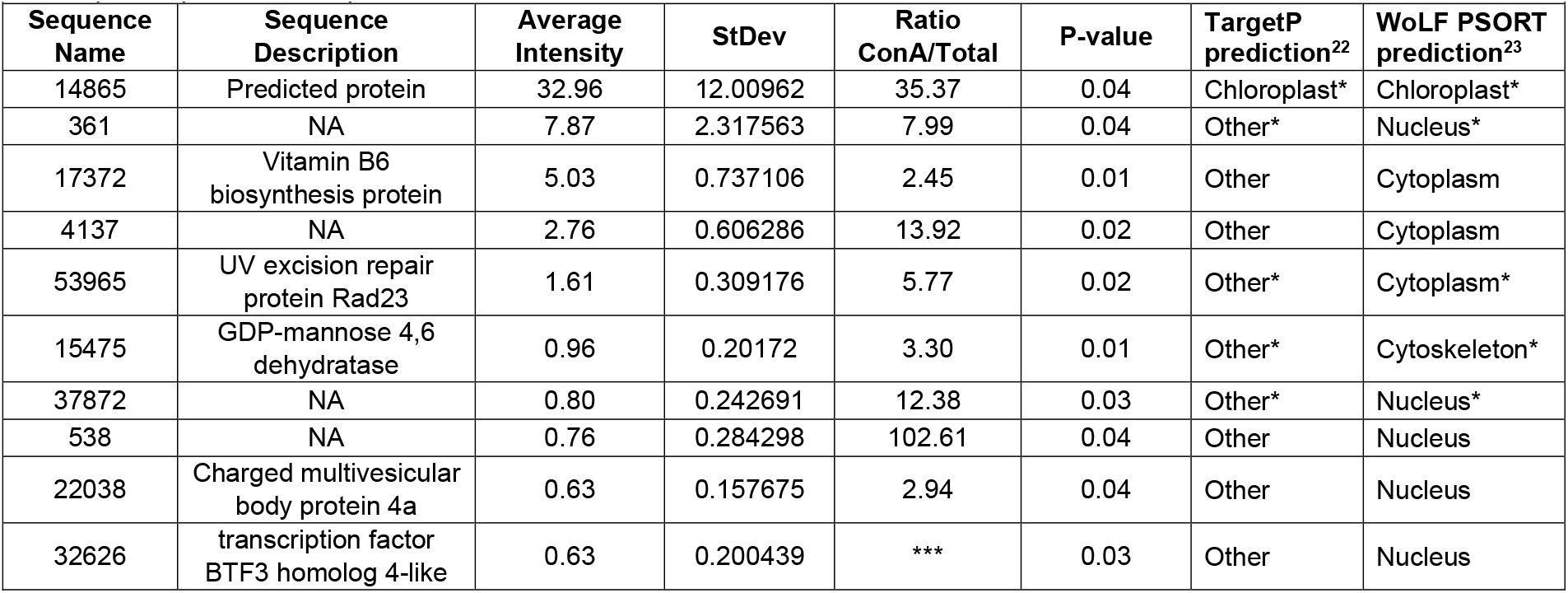
The 10 most abundant proteins enriched in ConA Glycoprotein Isolation. The full list is available in the supplementary file. NA is sequences with no homologues identified by BLAST. *** are proteins not detected in the total protein sample. * indicates sequences that do not start with a M and so may be truncated sequences that do not contain the targeting sequence present in the protein.

| Sequence Name | Sequence Description | Average Intensity | StDev | Ratio ConA/Total | P-value | TargetP prediction<sup>22</sup> | WoLF PSORT prediction<sup>23</sup> |
| --- | --- | --- | --- | --- | --- | --- | --- |
| 14865 | Predicted protein | 32.96 | 12.00962 | 35.37 | 0.04 | Chloroplast* | Chloroplast* |
| 361 | NA | 7.87 | 2.317563 | 7.99 | 0.04 | Other* | Nucleus* |
| 17372 | Vitamin B6 biosynthesis protein | 5.03 | 0.737106 | 2.45 | 0.01 | Other | Cytoplasm |
| 4137 | NA | 2.76 | 0.606286 | 13.92 | 0.02 | Other | Cytoplasm |
| 53965 | UV excision repair protein Rad23 | 1.61 | 0.309176 | 5.77 | 0.02 | Other* | Cytoplasm* |
| 15475 | GDP-mannose 4,6 dehydratase | 0.96 | 0.20172 | 3.30 | 0.01 | Other* | Cytoskeleton* |
| 37872 | NA | 0.80 | 0.242691 | 12.38 | 0.03 | Other* | Nucleus* |
| 538 | NA | 0.76 | 0.284298 | 102.61 | 0.04 | Other | Nucleus |
| 22038 | Charged multivesicular body protein 4a | 0.63 | 0.157675 | 2.94 | 0.04 | Other | Nucleus |
| 32626 | transcription factor BTF3 homolog 4-like | 0.63 | 0.200439 | *** | 0.03 | Other | Nucleus |

### 2.3 WGA Glycoprotein Isolation

Wheat Germ Agglutinin (WGA) is a protein that specifically binds GlcNAc (or sialic acid, which is not present in Euglena^16^), found in the core of *N*-glycans. A total of 675 proteins were detected in the sample eluted from the WGA Glycoprotein Isolation column. Of these, 16 were detected at a statistically significant rate higher than in the total cellular proteome (see Table 3), of which six were also detected in the ConA Glycoprotein Isolation sample. Just six of the 16 had matches in the non-redundant protein database and just four of these mapped to GO terms. These are a protein possibly involved in DNA repair, an oxidoreductase, a protein likely involved in retrograde signalling, and an integral membrane protease. It is possible that the WGA enriched proteins also contain an *O*-GlcNAc residue, a cytosolic protein modification found in eukaryotes with a role in cellular signalling and nutrient response.^24^

**Table 3:** Proteins enriched in WGA Glycoprotein Isolation. ^#^ Indicates proteins also identified as significantly enriched by ConA Glycoprotein Isolation. NA is sequences with no homologues identified by BLAST. *** are proteins not detected in the total protein sample. * indicates sequences that do not start with a M and so may be truncated sequences that do not contain the targeting sequence present in the protein. Any secretory signal peptides identified were removed and the analysis repeated, with results shown in brackets.

| Sequence Name | Sequence Description | Average Intensity | StDev | Ratio WGA/Total | P-value | TargetP prediction<sup>22</sup> | WoLF PSORT prediction<sup>23</sup> |
| --- | --- | --- | --- | --- | --- | --- | --- |
| 361<sup>#</sup> | NA | 5.612177 | 0.18 | 5.697825 | 0.000253 | Other* | Nucleus* |
| 24159 | UV excision repair protein RAD23 homolog B | 1.571933 | 0.41 | 5.800405 | 0.027599 | Other* | Chloroplast* |
| 9057<sup>#</sup> | NA | 0.481899 | 0.17 | *** | 0.040911 | Other | Nucleus |
| 20266<sup>#</sup> | S-(hydroxymethyl) glutathione dehydrogenase/class III alcohol dehydrogenase | 0.422701 | 0.12 | 6.008595 | 0.013053 | Other* | Cytoplasm* |
| 19374 | NA | 0.369053 | 0.13 | 22.50882 | 0.048467 | Other* | Nucleus* |
| 255 | Kinesin-like protein KIF16B | 0.187588 | 0.05 | 9.307156 | 0.026652 | Other | Cytoplasm |
| 59<sup>#</sup> | NA | 0.14351 | 0.05 | 4.492779 | 0.029378 | Other | Cytoplasm |
| 12064 | NA | 0.136116 | 0.04 | *** | 0.024536 | Other* | Cytoplasm* |
| 27863<sup>#</sup> | NA | 0.108907 | 0.00 | *** | 3.44E-05 | Other* | Mitochondria* |
| 53577 | NA | 0.104506 | 0.03 | *** | 0.023925 | Other* | Nucleus* |
| 5896 | NA | 0.055716 | 0.02 | 15.05286 | 0.045645 | Other* | Nucleus* |
| 17293 | NA | 0.05292 | 0.02 | *** | 0.03949 | Other | Chloroplast |
| 23739 | NA | 0.035889 | 0.01 | *** | 0.045216 | Other* | Nucleus* |
| 6278<sup>#</sup> | Predicted protein | 0.028572 | 0.01 | *** | 0.038147 | Other | Chloroplast |
| 8504 | NLR family CARD domain-containing protein 3-like | 0.010453 | 0.01 | 2.631579 | 0.044291 | Other* | Nucleus* |
| 4654 | S8 family serine peptidase | 0.007635 | 0.00 | *** | 0.013451 | Signal peptide (Other) | Chloroplast |

### 2.4 Extracellular proteome

As well as proteins isolated by lectin meditated enrichment, the extracellular proteome was analysed. These proteins were isolated from the cell-free media, and it should be noted that a small amount of extracellular media was included in the cell preparation for all other samples. A total of 135 proteins were detected in all three samples of the extra cellular proteome, of which 41 were not detected in the total proteome at all. 20 of these were statistically significantly more prevalent than in the total proteome (see Table 4), and of these only three had no BLAST matches (and only one further did not map to a GO term, despite matching a bacterial subtilisin related peptidase by BLAST). There are several proteins involved in transport and signalling. There is also a carbonic anhydrase, a thioredoxin, a peptidyl-prolyl cis-trans isomerase, a glycine dehydrogenase, and interestingly a possible protease inhibitor that could potentially be involved in pathogen resistance.^25^ There are also several proteins that would not be expected to be extracellular, such as a serine/threonine phosphatase, a chlorophyll binding protein and a CoA ligase. Interestingly the first and third most abundant proteins, also overrepresented in the ConA samples, do not match any sequences by BLAST.

**Table 4:** Proteins enriched in Extracellular proteome. NA is sequences with no homologues identified by BLAST. *** are proteins not detected in the total protein sample. Any secretory signal peptides were removed and the analysis repeated, with results shown in brackets. * indicates sequences that do not start with a M and so may be truncated sequences that do not contain the targeting sequence present in the protein.

| Sequence Name | Sequence Description | Average Intensity | StDev | Ratio Ext/Total | P-value | TargetP prediction<sup>22</sup> | WoLF PSORT prediction<sup>23</sup> |
| --- | --- | --- | --- | --- | --- | --- | --- |
| 2713 | NA | 72.80851 | 18.07437 | 20161.41 | 0.019932 | Signal peptide (Other) | Extracellular (Nuclear) |
| 11740 | S8 family peptidase | 0.570963 | 0.084867 | *** | 0.007284 | Signal peptide (Other) | Plasma membrane |
| 13218 | NA | 0.391458 | 0.058299 | 12426.41 | 0.007318 | Signal peptide (Other) | Extracellular (Cytosol) |
| 3168 | Long-chain-fatty-acid--CoA ligase 4 isoform X1 | 0.226524 | 0.080067 | 2644.55 | 0.039249 | Other* | Chloroplast* |
| 18460 | Hypothetical protein, conserved | 0.166714 | 0.062074 | *** | 0.043237 | Other* | Chloroplast* |
| 7055 | Light-harvesting chlorophyll a/b-binding protein | 0.14248 | 0.049539 | 45.55 | 0.039233 | Other* | Plasma membrane* |
| 1245 | P-ATPase family transporter | 0.114422 | 0.030738 | *** | 0.02322 | Other | Plasma membrane |
| 19656 | Cystatin B, putative | 0.095887 | 0.036113 | *** | 0.044171 | Other | Cytosol |
| 13679 | Serine/threonine-protein phosphatase PP-X isozyme 2 | 0.085083 | 0.022593 | 71.53 | 0.023225 | Other | Cytosol |
| 14610 | Carbonic anhydrase 1 | 0.061822 | 0.019603 | *** | 0.031917 | Other | Chloroplast |
| 6287 | CocE/NonD family hydrolase | 0.060295 | 0.024003 | 291.81 | 0.048503 | Other* | E.R.* |
| 1548 | aminomethyl-transferring glycine dehydrogenase | 0.058086 | 0.01343 | 12.89 | 0.023302 | Mitochondria | Mitochondria |
| 17754 | pleiotropic drug resistance protein ABC superfamily | 0.055424 | 0.012081 | 480.24 | 0.015448 | Other* | Nucleus* |
| 2873 | probable inorganic phosphate transporter 1-3 | 0.047583 | 0.013913 | *** | 0.027336 | Other* | Plasma membrane* |
| 19178 | 14-3-3 protein | 0.042734 | 0.003681 | 7.18 | 0.003449 | Other | Nucleus |
| 13675 | Putative 3, 5-cyclic nucleotide phosphodiesterase | 0.034218 | 0.0093 | *** | 0.023751 | Other | E.R. |
| 1168 | ATP-binding cassette sub-family G member 2 isoform X1 | 0.033678 | 0.00752 | *** | 0.016218 | Signal peptide (Other) | Plasma membrane |
| 34359 | Thioredoxin | 0.028584 | 0.007885 | 4.02 | 0.038769 | Mitochondria | Chloroplast |
| 26116 | Peptidyl-prolyl cis-trans isomerase CYP19-3 | 0.016344 | 0.003933 | 5.14 | 0.033596 | Mitochondria | Chloroplast |
| 27196 | NA | 0.015681 | 0.001283 | 130.58 | 0.002202 | Other* | Chloroplast* |

## 3. Discussion

As expected, the most abundant proteins in the total proteome were those associated with core housekeeping roles, central metabolism and the chloroplasts and mitochondria. Both ConA and WGA were able to enrich for a range of proteins, with some overlap, and the roles some of them may play on the cell surface can be postulated. The extracellular proteome has a number of proteins that could be involved in degrading extracellular material and signalling. The *N*-glycosylation site can be identified in some of the peptides, but it is notable that they are not reliably found at the canonical NX(S/T) sites of other Eukaryotes.

Of particular note are the large number of unique proteins, unrelated to any previously identified proteins, that are highly abundant in the total proteome, in the glycoprotein isolation samples and in the extracellular proteome. There are also several proteins which are only related to “predicted protein” and with no GO terms identified using Blast2GO. This data indicates there are a large number of highly abundant proteins in Euglena with no known function, some of which we can now tentatively identify as being glycosylated. These unique proteins would repay further study.

## 4. Methods

### 4.1 Culturing

*Euglena gracilis* Z (CCAP1224/5Z) was grown in 15 ml of EG:JM + glucose (15 g/L) at 30 °C with shaking (50 rpm) and illumination (800 lx) until late log phase (10 days) in triplicate. Cells were harvested by centrifugation (1000 xg) and resuspended in supernatant (1 ml).

### 4.2 Extra cellular proteins

The supernatant from the cell culture was filtered (0.2 µm) and lyophilised. The material was dissolved in ammonium bicarbonate (2.5 ml, 50 mM) and desalted using a PD 10 column (Amersham Pharmacia Biotech AG) equilibrated and eluted with ammonium bicarbonate (50 mM) the resultant material was again lyophilised and dissolved in MQ H_2_O (0.4 ml).

### 4.3 Glycoprotein preparation

The resuspended *Euglena* cells from the culturing (1 ml) were diluted with 5x Binding/Wash buffer (0.25 ml) containing phenylmethylsulfonyl fluoride (2 mM) and lysed by sonication (3 × 10 s, 25% amplitude, 30 s off between each pulse) and centrifuged (5 min, 1000 xg). Not all cells were lysed. Total lysate containing the equivalent of 1.1 mg of protein (Easy Bradford BioRad, BSA standards) was then used for glycoprotein purification using both ConA and WGA Glycoprotein Isolation Kits (Thermo Scientific) according to the manufacturer’s instructions. Protein quality was assed using silver stained SDS-PAGE (Bolt 4-12% Bis-TRIS plus, Invitrogen) using SeeBlue Plus2 Pre-stained Protein Standard (Thermo Fisher Scientific) as the standard.

### 4.4 Protein digestion and analysis by mass spectrometry

Protein digestion and analysis was performed by The Advanced Proteomics Facility at Oxford University. Protein samples were digested according to the Filter-Aided Sample Preparation (FASP) procedure.^26^ Peptide digest was treated with PNGase F and analysed by nano-liquid chromatography tandem mass spectrometry (nano-LC/MS/MS) on an Orbitrap Elite(tm) Hybrid Ion Trap-Orbitrap Mass Spectrometer (Thermo Scientific) using CID fragmentation. Peptides were loaded on a C18 PepMap100 pre-column (300 µm i.d. x 5 mm, 100Å, Thermo Fisher Scientific) at a flow rate of 12 μL/min in 100% buffer A (0.1% formic acid (FA) in water). Peptides were then transferred to an in-house packed analytical column heated at 45°C (50cm, 75 µm i.d. packed with ReproSil-Pur 120 C18-AQ, 1.9 µm, 120 Å, Dr. Maisch GmbH) and separated using a 60 min gradient from 8 to 30% buffer B (0.1% FA in acetonitrile (ACN)) at a flow rate of 200 nL/min. Survey scans were acquired at 120,000 resolution to a scan range from 350 to 1500 m/z. The mass spectrometer was operated in a data-dependent mode to automatically switch between MS and MS/MS. The 10 most intense precursor ions were submitted to Collision-Induced Dissociation fragmentation using a precursor isolation width set to 1.5 Da and a normalised collision energy of 35. Database search was carried out using MaxQuant (1.6.3.4) against the non-redundant Euglena proteome available at https://jicbio.nbi.ac.uk/euglena/,^12^ with default parameters and including Deamidation on Asn residues as variable modification for *N*-glycosylation sites identification. The mass spectrometry proteomics data have been deposited to the ProteomeXchange Consortium via the PRIDE^27^ partner repository with the dataset identifier PXD030579

### 4.5 Data Analysis

All Total, ConA and WGA samples were normalised with respect to an average of the 165 proteins detected in every sample. The Ext and Total samples were normalised to an average of the 81 proteins detected in both of these samples. Proteins that were differentially detected between the different treatments and the total proteome (P<0.05, Student’s T-test, two tail) were included in further analysis. Using Blast2GO, protein sequences were matched to sequences in the NCBI non-redundant protein sequence database and assigned GO terms based on this.

## 5. Supplementary Data

“Euglena glycoproteomics raw data” contains the raw data including the proteins, peptides, modifications and *N*-deamidation sites detected. “Euglena glycoproteomics data analysis” contains the data processed for all samples

## Supporting information

Euglena glycoproteomics raw data

Euglena glycoproteomics data analysis

## 6 Acknowledgments

This research was funded by a Glasstone Fellowship and a Nottingham Research Fellowship awarded to E. O’Neill. I would like to thank Rob Field, Eve Roxborough and Callum Southwood for helpful discussions of the manuscript.

